# A holistic insight of mycobacteriophage induced changes in mycobacterial cells

**DOI:** 10.1101/2021.10.21.465286

**Authors:** Fatema Calcuttawala, Rahul Shaw, Arpita Sarbajna, Moumita Dutta, Saptarshi Sinha, Sujoy K. Das Gupta

## Abstract

Mycobacteriophages are phages that interact with mycobacteria resulting in their killing. Although lysis is the major mechanism by which mycobacteriophages cause cell death, other mechanisms may also be involved. The present study was initiated with the objective of investigating the changes that take place at the cellular level following the infection of mycobacterial cells by phage D29. To investigate this issue, we took recourse to performing immunofluorescence and electron microscopic studies. Transmission electron microscopic examination revealed the adsorption of phages on to the surface of mycobacteria, following which penetration of the tail through the thick mycoloic acid layer was seen. At later time points discrete populations of cells at different stages of lysis were observed, which comprised of completely lysed cells, in which the cells were fragmented and those at the early onset stage exhibited formation of membrane pores through which the phages and intracellular contents were released. SEM results also indicated that phages may come out through the entire surface of the cell, or alternatively through gaps in the surface. In some of the images we observed structures that apparently resembled membrane blebs which are normally encountered when cells undergo programmed cell death (PCD). In addition, we observed significant increase in DNA fragmentation as well as membrane depolarization, which are also indicative of occurrence of PCD. As several bacterial PCD pathways are mediated by the toxin-antitoxin (TA) modules, the expression profile of all the TA systems was examined before and after phage infection. Apart from specifically addressing the issue of PCD in mycobacteriophage infected cells, this investigation has led to the development of facile tools necessary for investigating mycobacteriophage-mycobacteria interactions by means of microscopic methods.

## Introduction

Bacteriophages, literally meaning ‘bacteria devourers’, are the most abundant and diverse biological entities in the world (1). There are ~10^31^ phage particles in the biosphere (2). There is a dictum that phages are ubiquitously found where bacteria thrive, thereby playing a fundamental role in regulating bacterial ecology. They are obligate intracellular bacterial parasites, with either a lytic or a lysogenic life cycle (3). Even though they were discovered in 1915 by Frederick Twort, the nature of the existence of the so called “*contagium vivum fluidum*” and whether it was liquid or particulate remained a topic of contention until they were visualized for the first time in the year 1939 with an electron microscope (EM) by Knoll and Ruska (4,5). Several major milestones were achieved by phage imaging. These comprised of phage classification based on their morphological characteristics and studies on their interaction with the bacterial hosts (6).

The therapeutic potential of these transmissible bacteriolytic entities was first identified by their codiscoverer, Félix d’Hérelle (7). Since then phage research became the cradle of fundamental and translational biosciences. There is an increasing interest in the studies focusing on the use of bacteriophages as antibacterial agents against pathogenic bacteria. This is a consequence of the ability of the phage to lyse a bacterial host (8,9). Phage D29 is one such bacteriophage, which infects diverse mycobacteria such as *M. smegmatis* and *M. tuberculosis* (10). Belonging to the family Siphoviridae, it typically exhibits a long non-contractile tail (11). The resurgence of TB and emergence of excessive drug resistant (XDR) and totally drug resistant (TDR) strains has spurred renewed interest in the therapeutic use of mycobacteriophages (12). They can even serve as cornerstones for developing novel diagnostic and preventive strategies (13). D29 is the prototypical model for a mycobacteriophage as it efficiently adsorbs to the host and begins DNA replication within a few minutes after infection. One-step growth experiments have demonstrated that the length of the latent period, which is the time taken from infection to lysis, is 30-35 minutes in *M. smegmatis* but is 2-3 hours in *M. tuberculosis* (14,15). The rationale for the choice of *M. smegmatis* as the host is that it is non-pathogenic and grows substantially faster than *M. tuberculosis* (16). The kinetics of phage infection cycle is directly co-related to its host’s generation time and is extended in slow growing bacteria. For instance, DNA synthesis is observed 2-4 minutes after infection by *M. smegmatis* but is delayed to 20 minutes in case of the slow grower, *M. tuberculosis* (17). There are recent observations that *M. smegmatis* mc^2^155 is both restriction- and CRISPR-free suggesting that these are positive attributes for discovery of phages.

Preliminary studies in our laboratory have indicated that phage D29 infection results in ‘host inactivation’. However how this happens still remains obscure. Recent investigations have revealed that multiple mechanisms could be involved some of which are generation of superoxide radicals and induction of thymine less death (18,19). The present study was undertaken in order to gain an insight into the changes in the host cell upon phage infection. Electron microscopy, even though an age old technique, was resorted to, because it is still considered to be the gold standard for viral ultrastructure studies (20).

Several hurdles confront the utilization of phages for the curtailment of mycobacteria. Rather than recruiting phages directly for treatment, they can be used as platforms for drug discovery. Phages have evolved multiple strategies for interfering with bacterial growth. Understanding the targets that phages use in inhibiting bacterial growth has a clear therapeutic implication. In this study we focus on the interaction between mycobacteriophage and mycobacteria. Our objective was to develop cytological tools to understand the changes that happen within the bacterial cell once it is attacked by mycobacteriophages.

## Materials and Methods

### Bacteria, bacteriophage and media

Infection experiment was performed using *Mycobacterium smegmatis* mc^2^155 as the host strain and mycobacteriophage D29, which was obtained as a kind gift from Ruth McNerney (LSHTM Keppel Street, London, United Kingdom). Middlebrook 7H9 medium (Difco) supplemented with 0.2% glycerol, 0.25% bovine serum albumin (BSA) (HiMedia Laboratories, India) and 0.01% Tween 80 was used for growing mycobacterial cells. Phage infection was carried out in the same medium except that Tween 80 was omitted and 2 mM CaCl_2_ was added to the medium. MB7H9 hard agar plates were used for colony counting. The hard agar was overlaid with soft agar containing 2 mM CaCl_2_, for plaque assay.

### Phage infection assay

Phages were amplified by the confluent lysis method followed by suspension in SM buffer. Phage purification was done by performing CsCl density gradient centrifugation, followed by dialysis using a dialysis buffer (50mM Tris-Cl (pH 8.0), 10 mM NaCl, 10 mM MgCl_2_). *M. smegmatis* cells were infected with phage D29 at a multiplicity of infection (MOI) of 1. Aliquots were collected at different time points and centrifuged at 15,700X*g* for 5 min, the pellet and supernatant fractions were separated and the number of PFUs present in the pellet (infectious center) and the supernatant (free phage) were determined separately. The sum of the two values at time zero, immediately after phage addition, was considered as the input PFU. The MOI was determined by dividing the input PFU count by the total viable cell count, CFU, which was determined by plating the host cells on the same day.

### Cloning and expression of mycobacteriophage D29 gene *17*

The gene encoding the major head subunit gp17, was PCR amplified using the primers D2917F and D2917R (Table 1) from mycobacteriophage D29 genomic DNA and subsequently cloned into the BamHI-HindIII site of the expression vector pET-28a (Novagen). The recombinant protein had a His_6_ tag at the N-terminal end.

**Table 1.**
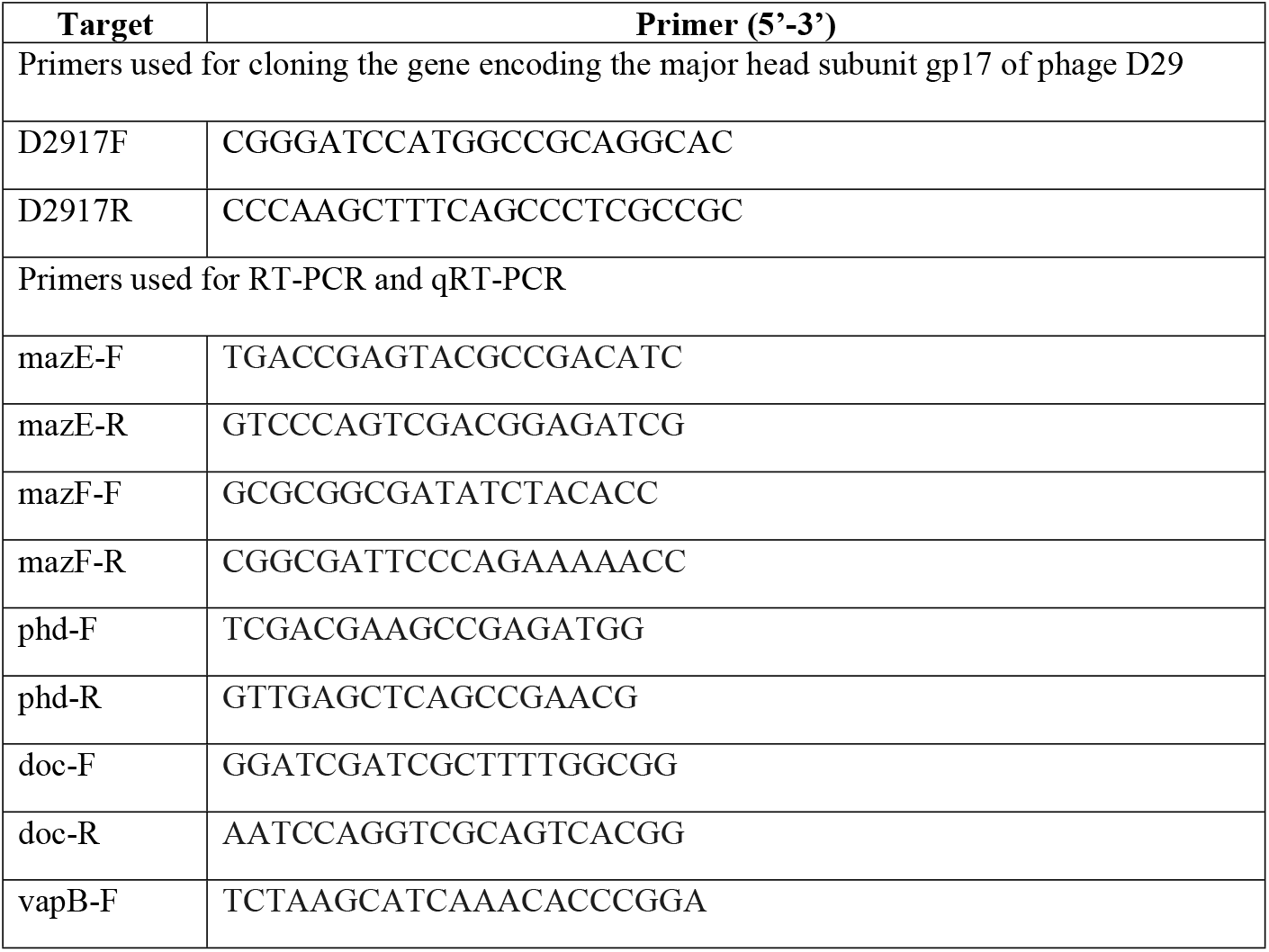
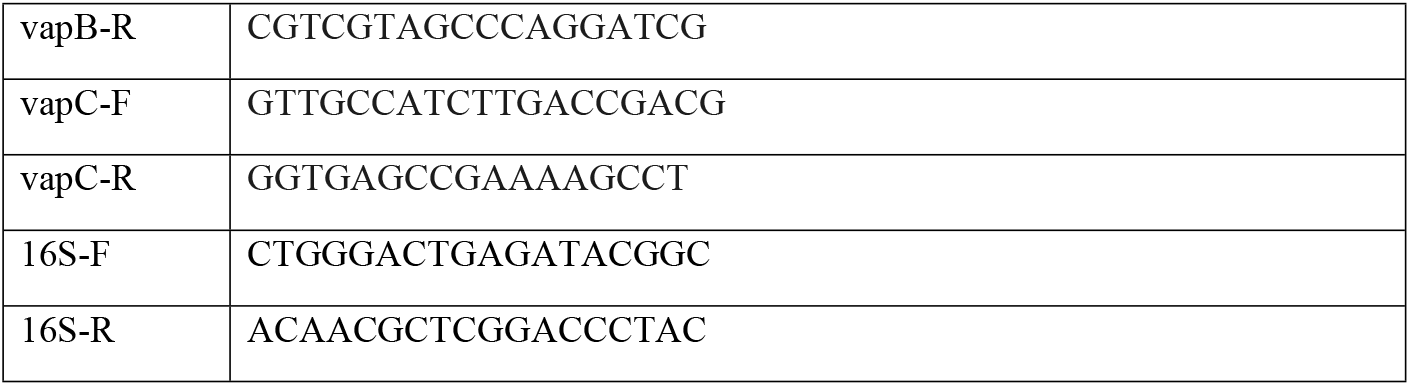
List of primers used in the study.

### Purification of recombinant protein

The plasmid construct made for over-expressing the protein gp17 was transformed into *E. coli* BL21(DE3). The transformants resistant to kanamycin (50μg/ml) were cultured in the presence of the antibiotic at 37°C. At an OD_600_ of 0.5, isopropyl-β-D-thiogalactopyranoside (IPTG) was added at a final concentration of 0.5 mM. The bacterial cells were induced at 37°C for 3 hrs. Bacterial pellets were obtained by centrifugation at 10,000X*g* for 15 min. Cells harvested by centrifugation were lysed by sonication. Protein purification was performed using Ni^+2^-NTA agarose affinity chromatography according to standard protocol (Qiagen).

### Raising antibody in rabbit

Polyclonal antibodies were raised against affinity-purified gp17 protein which was isolated under denaturing conditions in the presence of 8 M urea. The protein sample was gel purified and injected into rabbit. Pre-immune and immune sera were drawn and the specificities of the sera thus obtained were verified by Western blotting.

### Immunofluorescence microscopy studies

At both early and late stages of infection, cells were harvested, washed with PBS and then fixed in 4% (w/v) paraformaldehyde in PBS for 20 min at room temperature. After several PBS washes, blocking was performed using the blocking buffer (3% BSA in PBS) for 15 min. Cells were subsequently treated with primary antibodies in blocking buffer against gp17 at 4°C for 1 hr. After several washes with PBS, the cells were incubated with Dylight488-labelled goat anti-rabbit secondary antibodies (Thermo Fisher Scientific, USA) in blocking buffer at 4°C for 1 hr. After several washes with PBS, cells were stained with 4’, 6’-diamidino-2-phenylindole (DAPI) to visualize the nucleic acid. Stained cells were examined by confocal microscopy (Leica TCS SP8).

### Transmission Electron Microscopy (TEM) studies

The interaction between the mycobacteriophage D29 and its host strain was examined by transmission electron microscopy. Infection was performed at an MOI of 1. Samples collected at different stages of infection were negatively stained with 2% uranyl acetate and examined under a FEI Tecnai 12 Biotwin transmission electron microscope (FEI,Hillsboro,OR,USA) at an accelerating voltage of 100kV.

### Scanning Electron Microscopy (SEM) studies

For scanning electron microscopy, samples were collected and fixed with glutaraldehyde (1.5% w/v). After centrifugation at 3,300X*g*, the pellet was dissolved in 20% ethanol solution and spread on the glass slide for drying, mounted on aluminium stubs, coated with gold (Edwards) and photographed with a SEM (FEI Quanta 200).

### Membrane depolarization assay by flow cytometry

Cultures were grown overnight at 37°C to an OD_600_ between 0.2-0.3 before phage treatment. Infection was done at an MOI of 1. After treatment the sample was collected, washed once and resuspended in 1 ml 1X PBS. Staining of cells was performed using 5 μl of DiBAC_4_(3) (Invitrogen, 0.025 mg/ml in DMSO) followed by incubation for 15 minutes at room temperature in dark. The intensity of DiBAC_4_ fluorescence was measured using the FACSVerse (BD) system with a 488-nm argon laser for excitation and a 530±15-nm emission filter.

### DNA fragmentation assay by flow cytometry and confocal microscopy

DNA fragmentation was quantified by the TUNEL assay using the ApoDirect kit (BD Biosciences). The enzyme terminal deoxynucleotidyl transferase (TdT) adds fluorescein isothiocyanate deoxyuridine triphosphate (FITC-dUTP) to each 3’-hydroxyl end of fragmented DNA, making it possible to determine the extent of DNA fragmentation by measuring the intensity of fluorescence. For our studies, cells were grown and treated with phage. After washing with PBS, cells were fixed by resuspending them in 1 ml of 1% paraformaldehyde and incubating on ice for 60 min, following which they were washed with PBS, resuspended in 70% ethanol and stored at −20°C overnight. On the next day, the cells were centrifuged, ethanol was discarded and the cell pellet was resuspended in 1 ml wash buffer provided in the kit. After two washes using this buffer, the pellet was resuspended in 50 μl of a staining solution that comprised of reaction buffer, FITC-dUTP and TdT dissolved in distilled water. The reaction mixture was incubated at 37°C for 60 min, gently mixing the sample after every 15 min. The reaction was arrested by adding 1 ml of rinse buffer from the kit. The rinse was performed twice followed by addition of 500 μl prodium iodide/RNase solution and incubation in the dark for 30 min at room temperature. The PI/RNase treatment reduces RNA-related background, and also reports the DNA content of the labelled cells. The intensity of fluorescence was measured using the FACSVerse (BD) system. An aliquot of the stained cells was centrifuged, resuspended in 1X PBS and examined by confocal microscopy. (Leica TCS SP8).

### RNA extraction, RT-PCR and qRT-PCR

RNA was extracted from 20 ml of broth culture. Cells were harvested by centrifugation (13,000X*g*, 5min) and re-suspended in 2 ml TE buffer (10 mM Tris-HCl pH 8, 1 mM EDTA) containing lysozyme at a concentration of 5 mg/ml, followed by incubation at 37°C for 30 min. This was followed by addition of 5 ml TRIzol reagent (Invitrogen), 1ml chloroform and centrifugation at 13,000X*g* for 15 min. The aqueous phase was collected and precipitated by adding 0.7 volumes of cold isopropanol, followed by 70% ethanol wash. The pellet was dried and re-suspended in 50 μl of DEPC treated water. Contaminating DNA was removed by DNaseI digestion (Qiagen) and RNA clean up (RNeasy Mini Kit, Qiagen) performed according to the manufacturer’s instructions. RNA quality was assessed by agarose gel electrophoresis and A_260/280_.

First strand cDNA synthesis was done from 400 ng RNA using RevertAid Reverse Transcriptase (Fermentas) by random priming according to the manufacturer’s instructions. One microliter of cDNA, 0.25mM dNTPs, 0.2 pmol/ μl of each primer and 0.3U of Taq polymerase were used for 25 μl of RT-PCR reaction mixtures. Thermal cycler conditions were: 94°C for 5 min, 30 cycles of 94°C for 30 sec, 55°C for 30 sec followed by 72°C for 7 min and a final hold at 4°C. 16S rRNA transcript levels were used as endogenous control. The PCR products were analysed by agarose gel electrophoresis.

Real-time analysis was conducted using Power SYBER^R^ Green PCR Master Mix (Applied Biosystems) in a 7500 Fast Real-Time PCR system (Applied Biosystems). The cycling conditions were: 50°C for 2 min, 95°C for 30 sec, 55°C for 30 sec, 72°C for 45 sec. All the primers used for RT-PCR and qRT-PCR are enlisted in Table 1.

## Results

### Immunofluorescence microscopic studies of phage host interaction

The interaction of mycobacteriophage D29 with its host bacterium *M. smegmatis* mc^2^155 was visually inspected by means of immunofluorescence microscopy. Antibodies were raised in rabbit against the phage D29 capsid protein gp17. Cells were subjected to immunofluorescence staining using anti-rabbit Dylight488 labelled secondary antibody. Pre-immune serum was found to contain no gp17 specific antibodies, as confirmed by Western blot and confocal microscopy. Our results revealed distinct phage adsorption 30 min post-infection. After 2 hrs, several free phages were observed, indicative of phage release after lysis (Fig 1).

**Fig 1.**
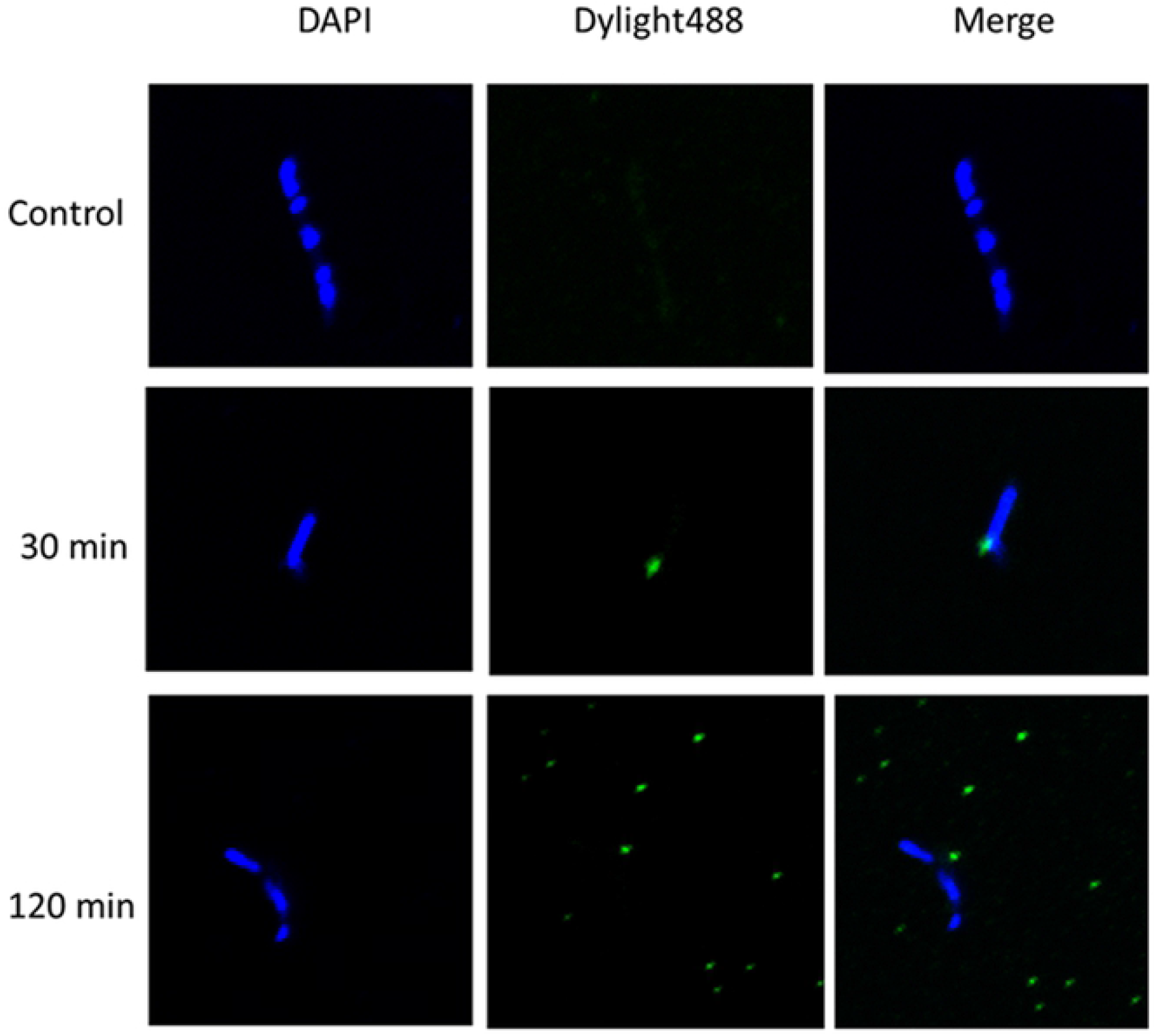
Immunofluorescence microscopy of control and phage infected cells. Phages were detected by staining with polyclonal rabbit serum against gp17 capsid protein, followed by anti-rabbit Dylight488 conjugated secondary antibody and cells were localized by DAPI, which stains the nucleic acid.

### Phage induced morphological changes in mycobacteria by electron microscopy

Morphological examination of the phage D29 revealed that it had an icosahedral head with a diameter of ~50 nm and a ~100 nm long tail, at the distal end of which tail fibers were occasionally observed (Fig 2A). The mycobacterial cells measured ~3-4 μm lengthwise and exhibited a typical thick cell envelope (Fig 2B). During the early stages i.e., 30 mins post infection, the adsorption of the phage particles to the host cell was clearly observed. Even though infection was performed at an MOI of 1, multiple phages were seen adsorbed to the bacterial cell surface (Figs 3A, B, E, F). Phages were attached both at the extremities as well as a localized portion on the surface of the mycobacterial cell. Following phage adsorption, penetration of the tail through the thick mycolic acid layer was observed (Figs 3C and 3D). Once the tail pierced the cell wall, the phages retained their normal appearance for a very short duration of time. Post-DNA injection, “ghost” phage particles, which are empty capsids were occasionally found attached to the surface (Figs 3E and 3F). Distinct protrusions of the membrane, analogous to membrane blebs were also observed (Figs 4A-C). At a later stage, i.e., 2 hrs post infection, several discrete populations of cells were observed. Some cells exhibited several pores in the membrane (Figs 5A-F). Leakage of intracellular contents and release of phages was observed from these pores. Several mature phages were seen clustered and localized close to the host cell (Figs 5C and 5D). These cells may exhibit the onset of lysis. Another population consisting mainly of fragmented cells and cellular debris was seen, representing later stages of lysis (Fig 6). SEM data correlated well with that of TEM (Fig 7).

**Fig 2.**
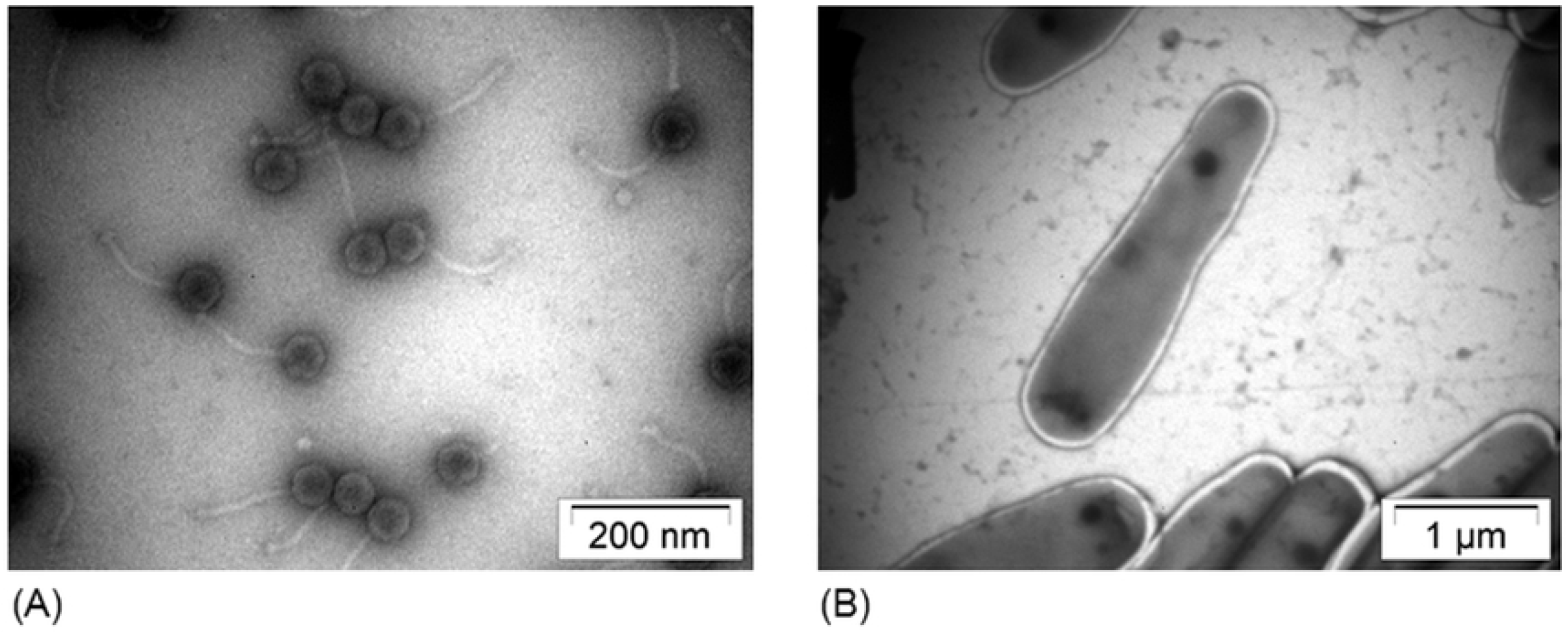
TEM image of (A) phage D29 (B) *M.smegmatis* mc^2^155 cells.

**Fig 3.**
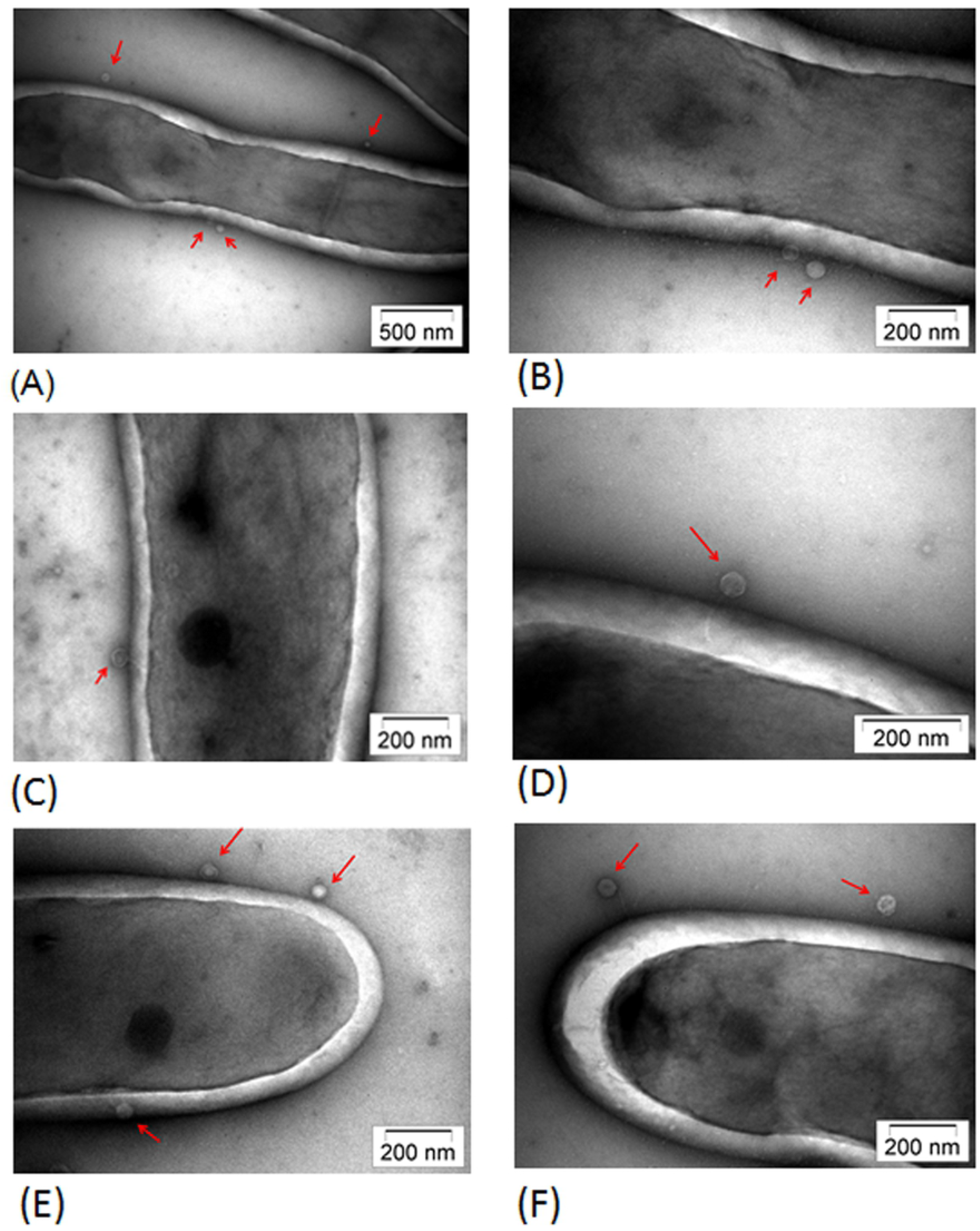
TEM images showing phage adsorption. Red arrows denote phages.

**Fig 4.**
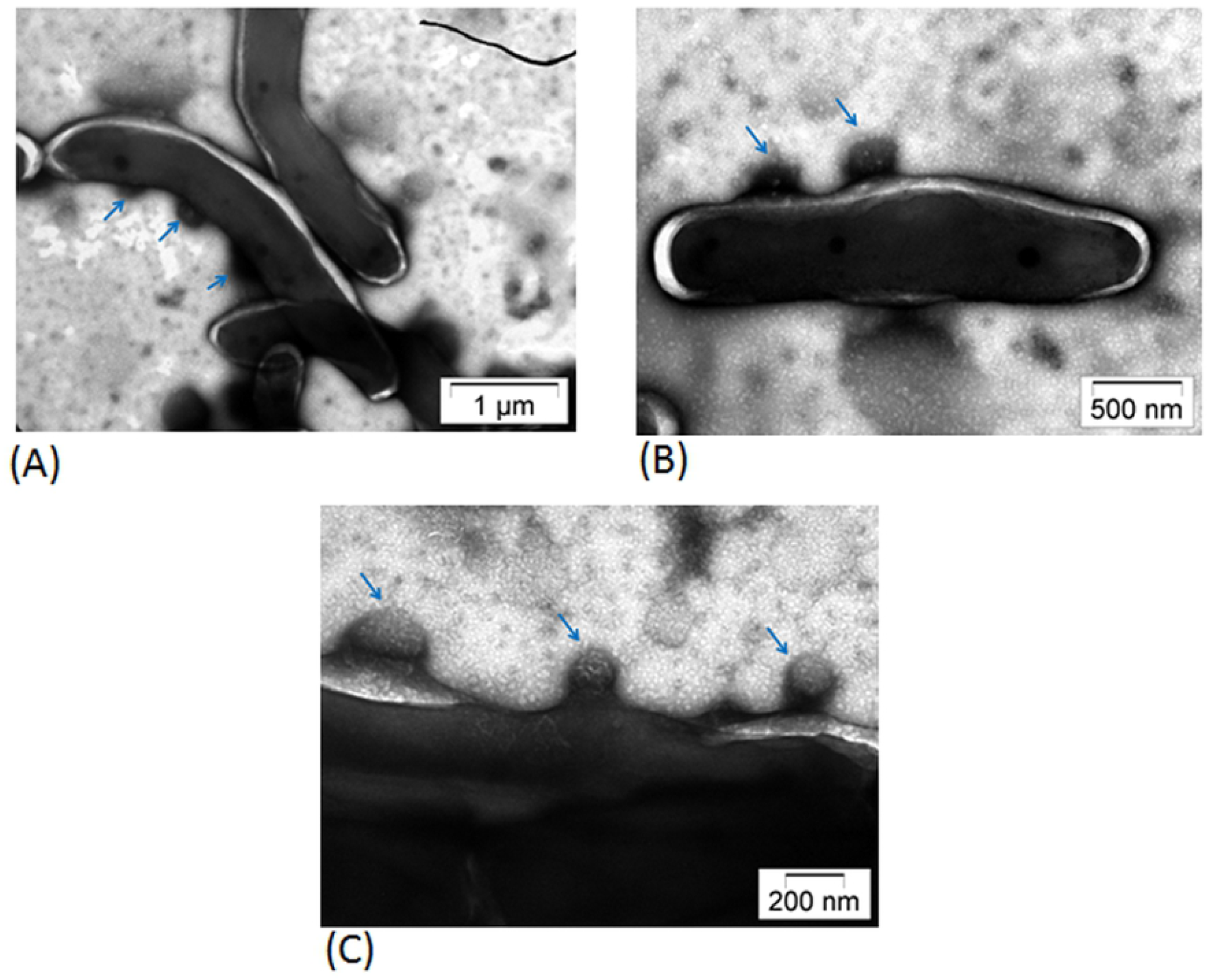
TEM images showing membrane blebbing during early stages of phage infection. Blue arrows denote membrane blebs.

**Fig 5.**
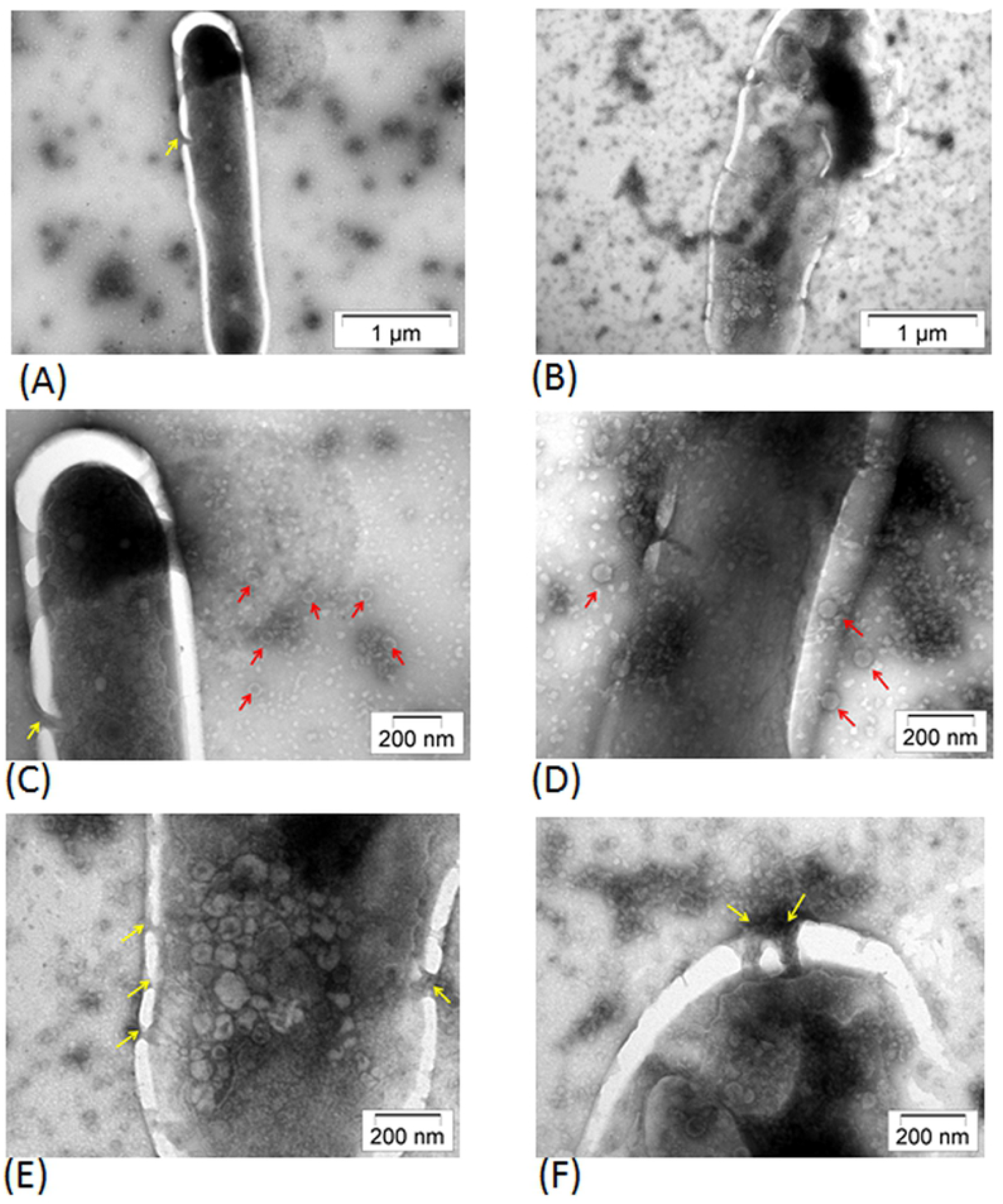
TEM images showing cellular changes during early onset of lysis (red arrows denote phages and yellow arrows indicate pore formation in the membrane).

**Fig 6.**
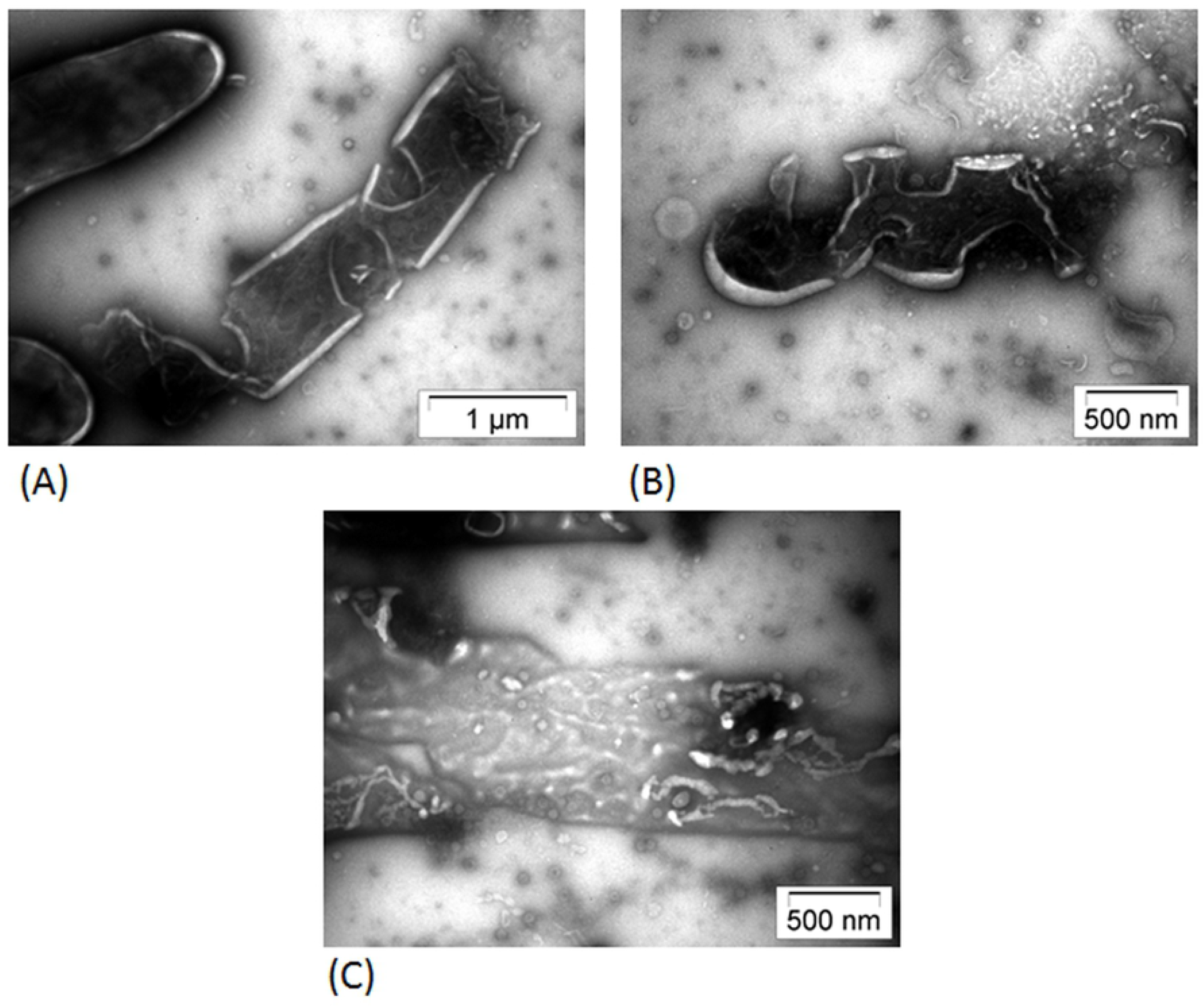
TEM images showing cellular changes during late stages of lysis.

**Fig 7.**
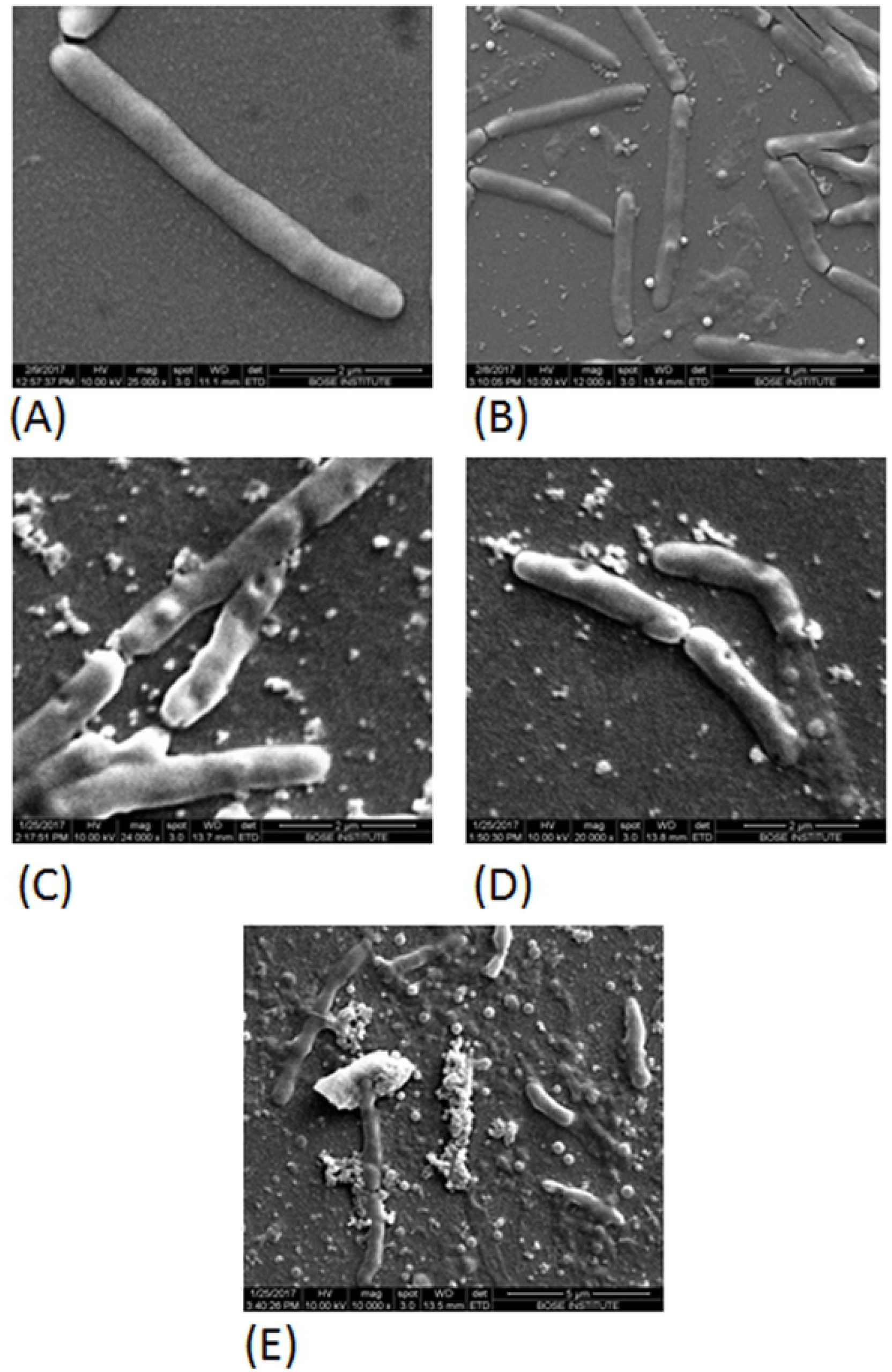
SEM micrographs depicting (A) *M. smegmatis* mc^2^155 (B) Phage adsorption (C) Membrane blebbing (D) Pore formation in the membrane (E) cell lysis.

### Phage infection induces membrane depolarization

Membrane depolarization is routinely monitored using the membrane potential sensitive dye bis-1,3-dibutylbarbituric acid trimethine oxonol (DiBAC_4_), which diffuses across depolarized yet intact cell membranes, and binds to lipid rich intracellular components. We used DiBAC_4_ staining together with flow cytometry in order to detect fluorescence in individual dying cells. Polymyxin B (50 μg/ml) treated cells were included as positive control. To our surprise, even the untreated cells exhibited some basal fluorescence. This was later found to be due to the addition of Tween to the culture at the time of inoculation. However, upon 4 hours of phage treatment, the level of DiBAC_4_ fluorescence increased substantially (~4 fold increase compared to the control set), indicating a loss of membrane potential (Fig 8).

**Fig 8.**
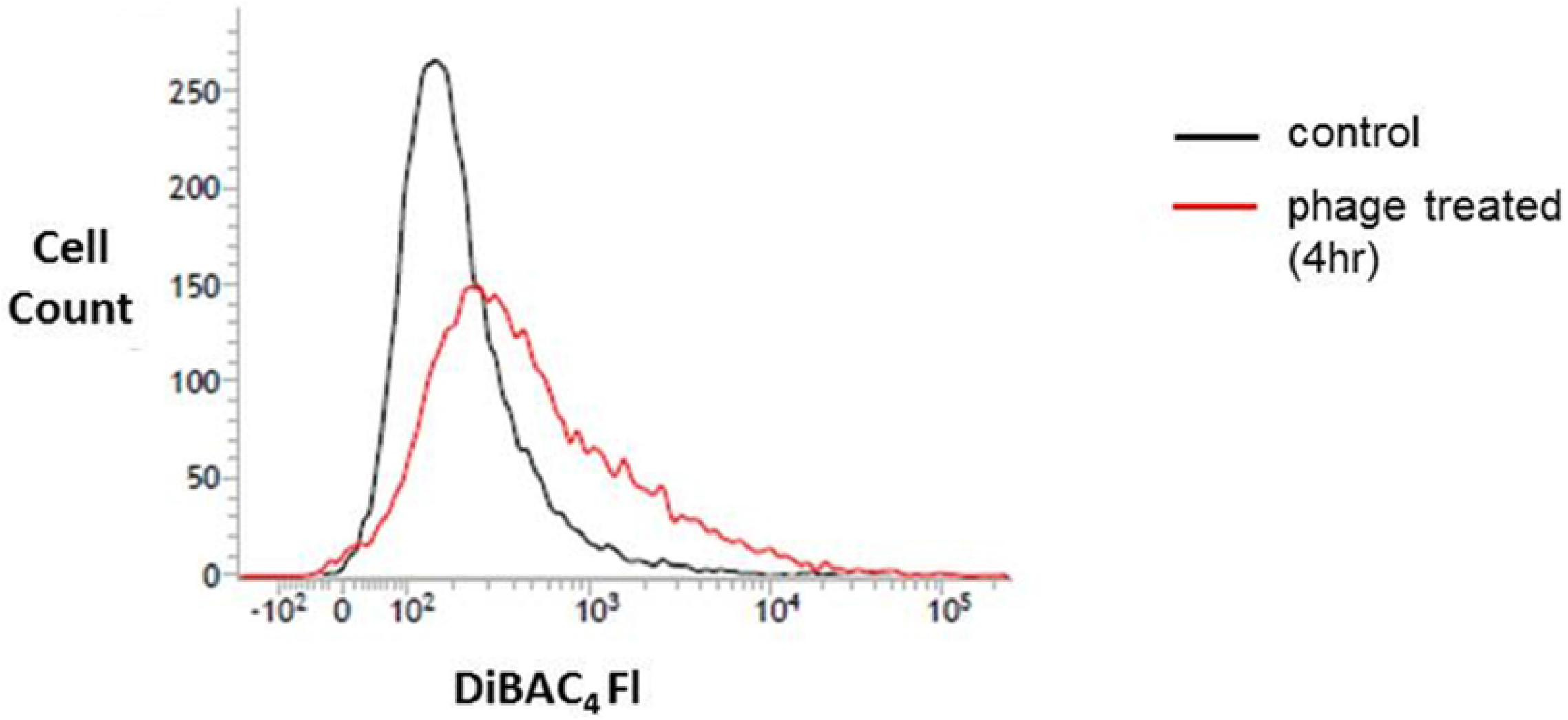
DiBAC_4_ (3) staining as a measure of membrane depolarization. Fluorescence intensity was measured by flow cytometry. This data represents the results of one of the three similar experiments.

### Phage infection induces TUNEL-detectable DNA fragmentation

DNA fragmentation was evaluated by the dominant method of TUNEL assay using the APO-Direct kit. The fluorescence of individual cells was measured by flow cytometry. Mycobacterial cells treated with hydroxyurea (20 mM) for 8 hours were used as positive control. In case of phage treated cells, no appreciable DNA fragmentation was observed at the early stages of infection. However, an increase in the percentage of TUNEL positive cells (~3 fold increase compared to the control set) indicating an increase in DNA damage was observed after 4 hours of infection (Fig 9). Flow cytometry data was further supported by direct microscopic assessment using PI and FITC to stain the *M. smegmatis* mc^2^155 cells. Phage treated cells stained positively for PI and FITC as compared to control cells which were stained with PI only (Fig 10).

**Fig 9.**
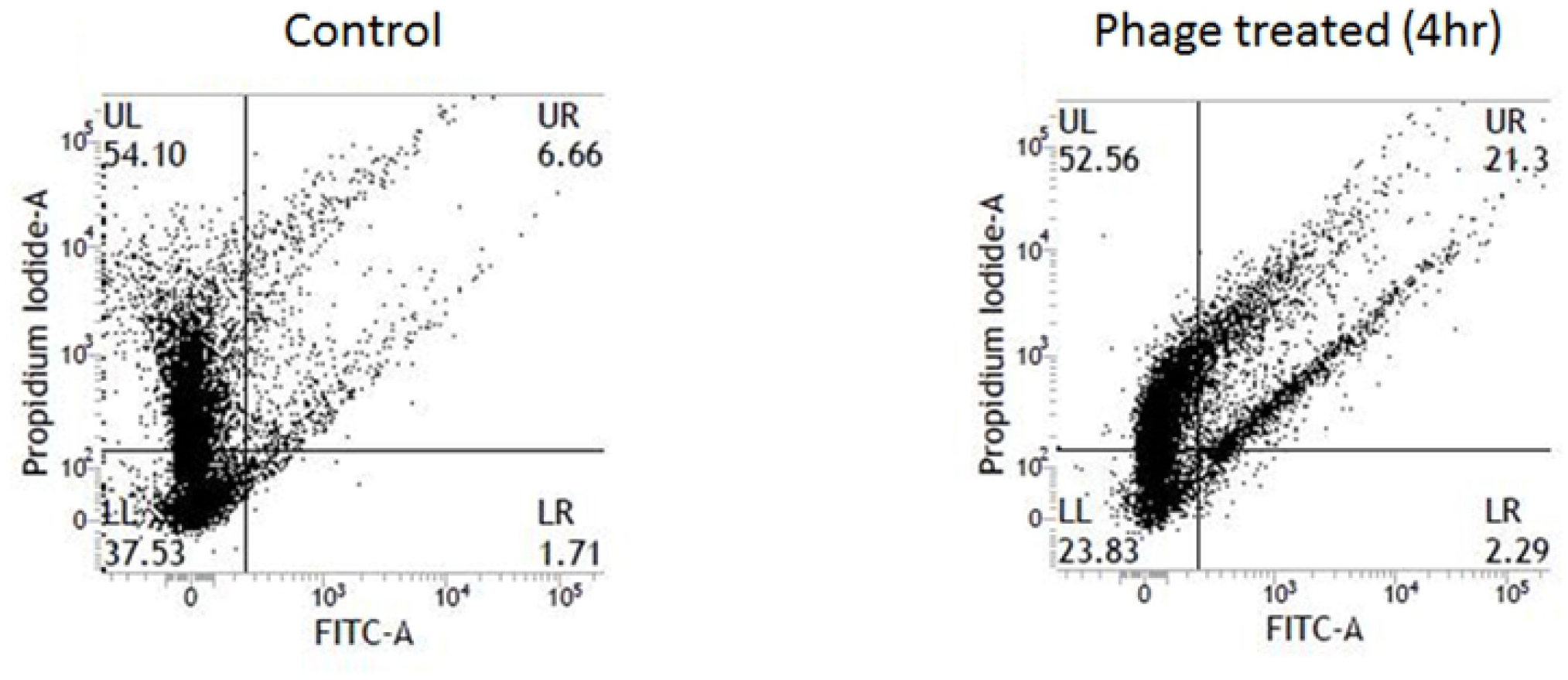
DNA fragmentation detected by the TUNEL assay. UL (Upper Left), UR (Upper Right), LL (Lower Left) and LR (Lower Right) denote the percentage of cells in each of the quadrants. The threshold was set using untreated and unstained *M.smegmatis* mc^2^155 cells. This data represents the results of one of the three similar experiments. FITC-fluorescein isothiocyanate, SSC-side scatter.

**Fig 10.**
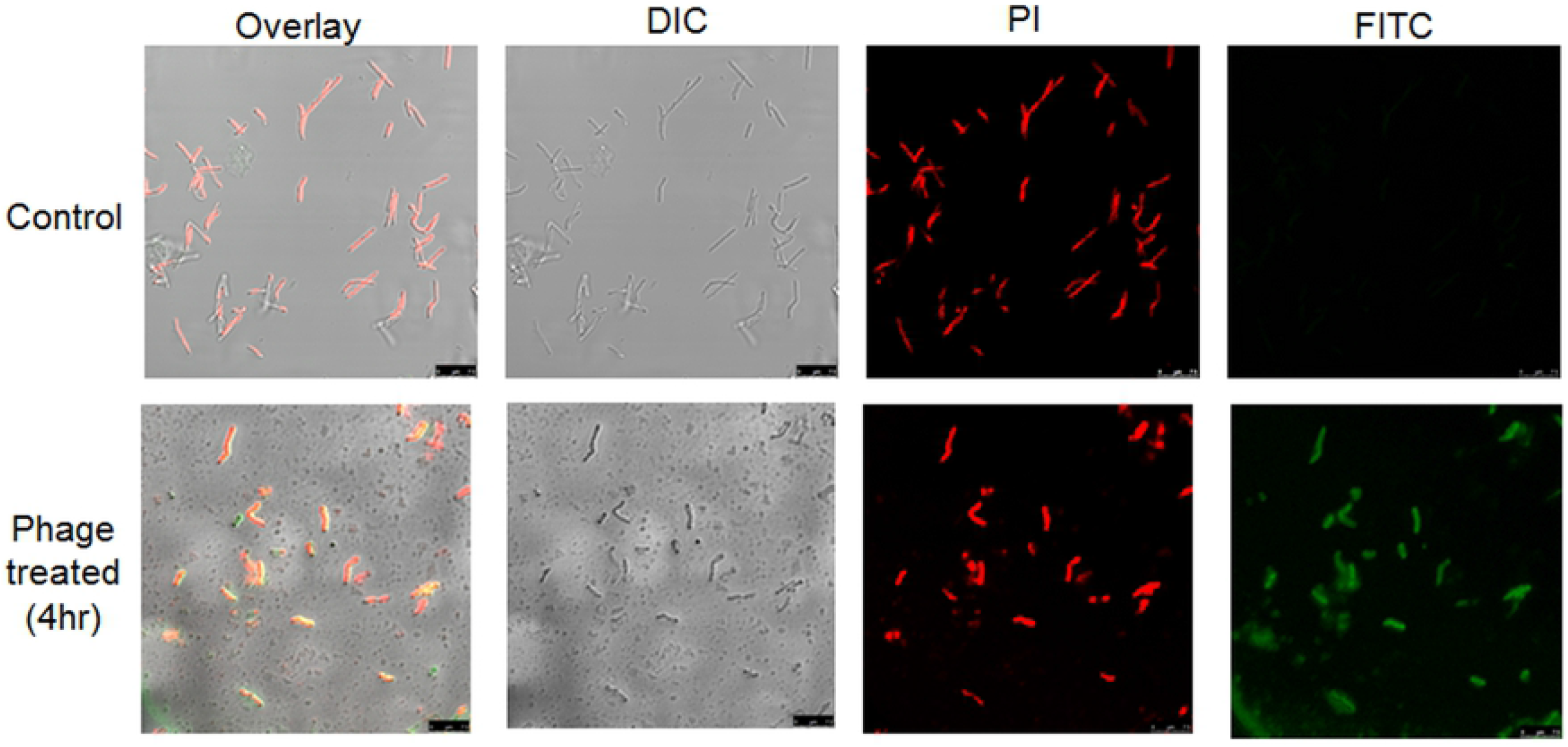
Microscopic analysis of phage infected and control cells stained with the dyes PI and FITC. Scale bars 7.5 μm.

### Expression profiling of toxin-antitoxin systems of the host upon phage infection

*Mycobacterium smegmatis* contains three putative toxin-antitoxin (TA) systems MazEF, VapBC and Phd/Doc (21). TA modules comprise of a pair of genes usually co-transcribed as an operon, in which the downstream gene encodes the stable toxin and the upstream one encodes the labile antitoxin (22). In our study, the expression profile of all the TA systems was analyzed at two time points, 2 hrs and 4 hrs respectively, upon phage infection. A unified pattern of expression was observed. The RT-PCR results indicated that in general the expression level of all the TA system genes decreased from 2 hrs to 4 hrs (Fig 11) in the phage infected sets. To further ensure that these changes were due to phage infection, PCR was performed using primers for the phage D29 gene encoding gp88 (data not shown). In case of all the samples, no amplification products were observed in the absence of reverse transcriptase, thereby confirming that the products seen by RT-PCR were due to the TA transcripts and not DNA contamination. Quantitative measurement by real-time PCR was done for the VapBC system and the qRT-PCR results were found to be consistent with those obtained by the qualitative end point PCR. A dramatic decrease in gene expression was observed upon phage infection (Fig 12) whereas the expression was found to increase marginally in the uninfected controls.

**Fig 11.**
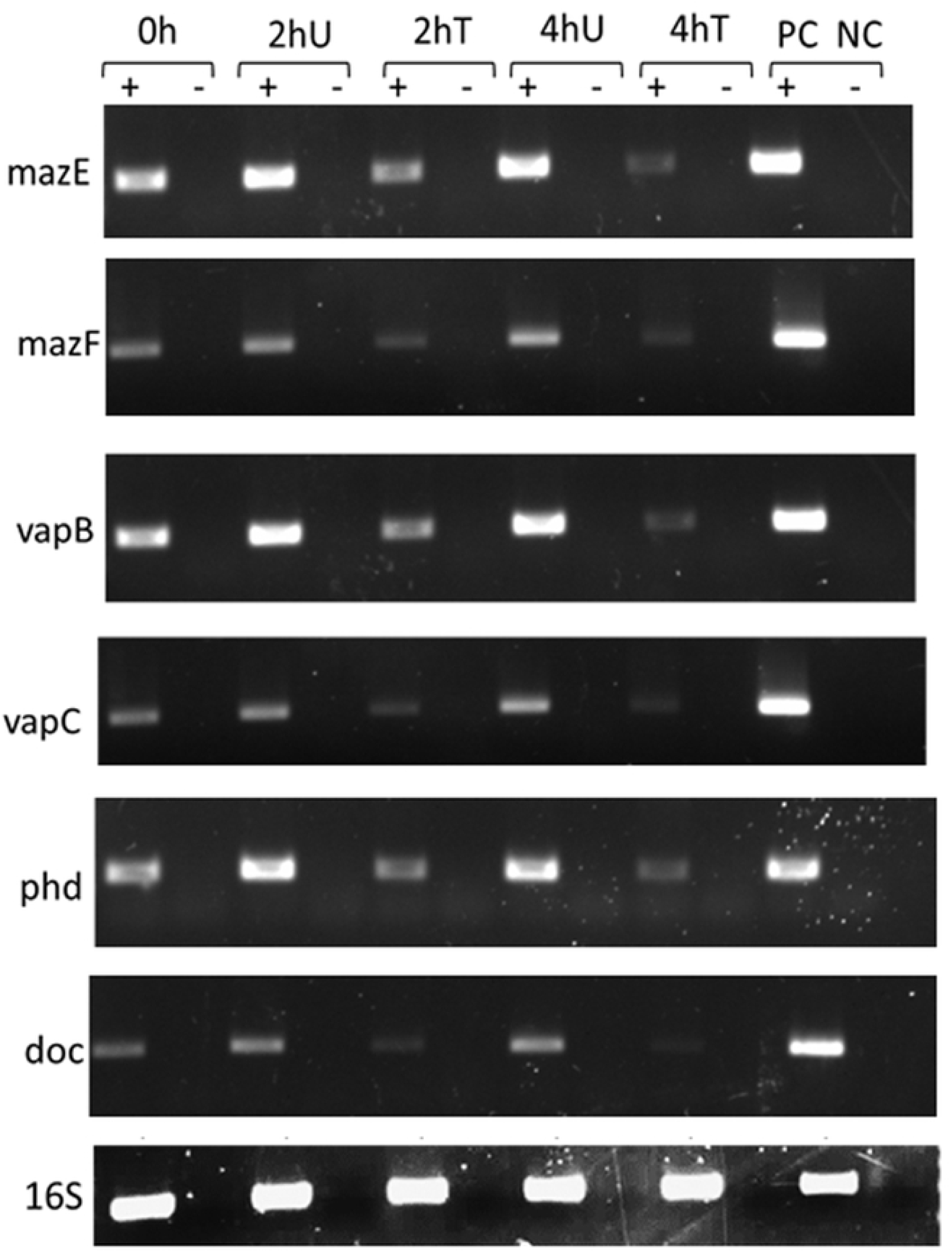
RT-PCR analysis of the various toxin-antitoxin genes of *M.smegmatis* mc^2^155. In each case, the name of the target gene is mentioned on the left side of the panel and the time points at which RNA was extracted from the cells are mentioned on the top. To ensure that the bands represent RNA entities and not their DNA counterparts, RT-PCR analysis was performed with samples that were either subjected (+) or not subjected (-) to reverse transcription prior to PCR. As internal control, amplification was done using 16S rRNA primers. U-Untreated, T-D29 phage treated, PC-Positive control, NC-Negative control.

**Fig 12.**
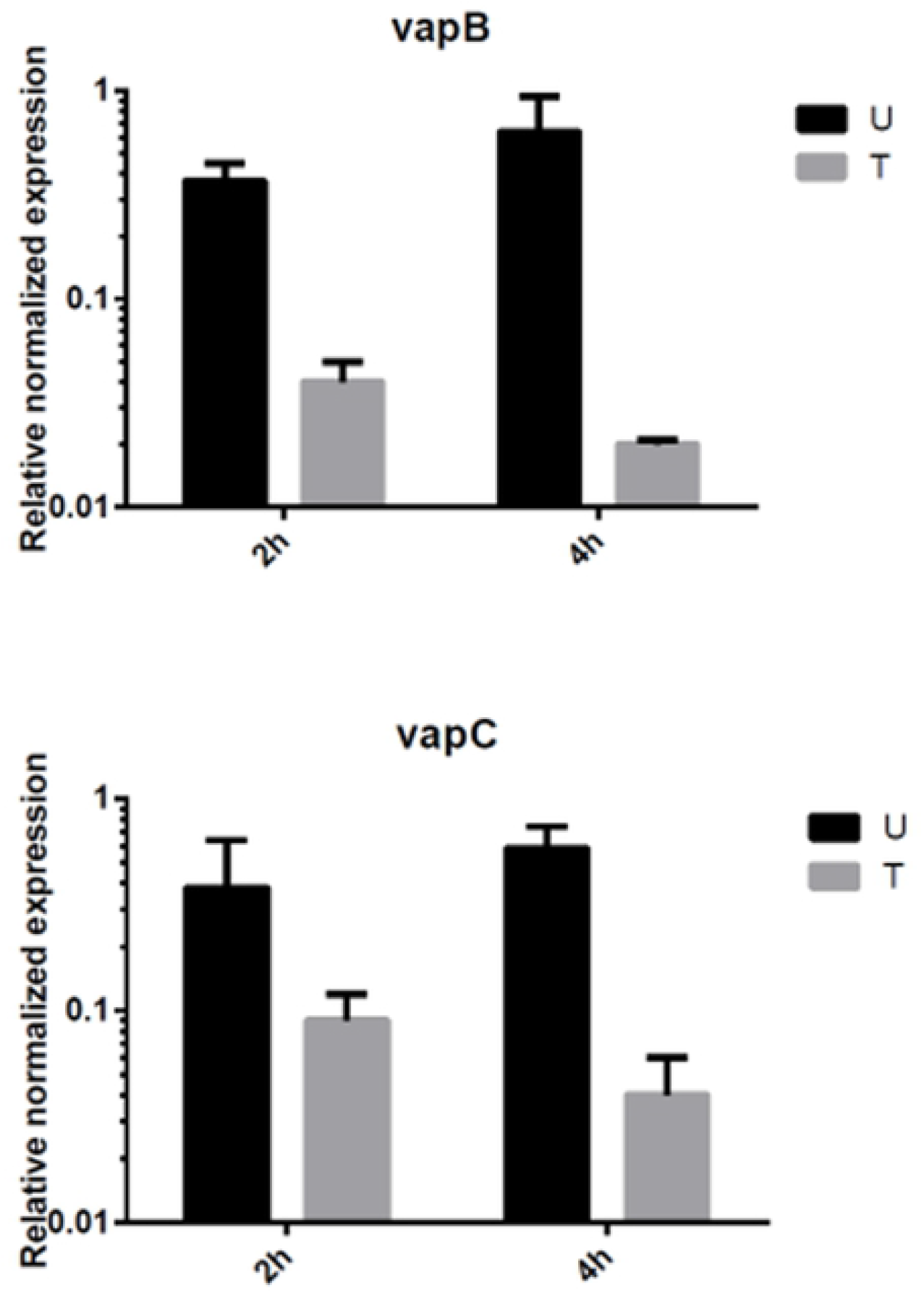
Analysis of VapBC gene expression 2 hrs and 4 hrs after phage D29 infection by qRT-PCR. The expression level of each gene was determined by the comparative C_T_ method after normalizing with a 16S rRNA control. The data were averaged from three independent experiments ± S.D.

## Discussion

The present investigation was undertaken in order to determine the morphological and physiological changes that mycobacterial cells undergo upon phage infection. Past speculation reveals that for a long time the research with mycobacteriophages possessed a merely phenomenological character, where it was possible to observe productive infection by the phages only by scoring for the efficiency of plaque formation (23). However, with the advent of immunofluorescence microscopy, the ‘fluorophages’ provided a visual alternative for mechanistically dissecting the crucial steps of phage adsorption and lysis, leading to plaque formation (24,25). In this work we examined phage-host interaction at the single-phage, single-cell level and thereby present a new tool for studying different properties of infection propagation at the microscopic level by employing a ‘fluorophage’. Even though our work is at its infancy, yet it provides a stepping stone towards performing time lapse experiments under the microscope. This in turn would provide a more precise understanding of the dynamics of infection and render complete knowledge regarding the propagation of infection in real time.

Electron microscopy took our studies on understanding phage-host interaction to a fundamentally new level (26). Right from the past, the mycobacteria-mycobacteriophage system has been considered to be ideal for EM studies because the two-fold obstacle confronted using *E. coli* and their phages could be overcome by using this model system (27). The obstacles comprised of opacity of the *E. coli* cell to electrons, which hindered the observation of the cell interior and the extreme rapidity of lysis, which made it impossible to distinguish between different phases of the infection process (28). However, the electron transparency of mycobacterial cells upon phage attack and prolonged infection cycle of mycobacteriophages acted as positive attributes for EM studies (29). This has made it possible for us to visualize the complete infection process from the beginning till the end, including phage adsorption, penetration of the non-contractile tail through the thick mycolic acid layer, the morphological changes in the infected cells including pore formation during onset of lysis, followed by bursting of the host cells and release of mature phages and intracellular contents. Non-contractile tails are a defining feature of phages belonging to the Siphoviridae family, of which phage D29 is a member (11). Another striking observation was that the dimensions of the pores in the membrane were intriguingly smaller than the phages emerging from them. Also, the membrane of the infected mycobacterial cells showed structures resembling membrane blebs, which is a characteristic feature of cells undergoing PCD (30).

PCD is a genetically-determined process characterized by a stereotypical set of morphological hallmarks (31,32). Our results demonstrated that phage infection induced characteristic changes in the host cells, such as DNA fragmentation, membrane depolarization and membrane blebbing that are strikingly reminiscent of eukaryotic apoptosis. Depolarization of the mitochondrial inner membrane is one of the key events which characterize the intrinsic pathway of apoptosis in case of eukaryotes (33). The phage D29 holin-like proteins and the eukaryotic BCL-2 family of proteins seem to utilize similar strategies to disrupt the integrity of the host cell membrane (34). Bacteria are ancestral to mitochondria as evidenced by the endosymbiotic theory (35,36). Thus, taking these similarities into consideration, it is tempting to speculate the possibility of convergent evolution of the machinery that controls PCD in case of prokaryotes and eukaryotes.

Previously, experiments conducted in our laboratory had demonstrated that the decrease in number of viable cells far exceeded the number of cells that undergo lysis upon phage infection. This phenomenon of death without lysis (DWL) was proposed to involve superoxide radical generation but other secondary factors remained unexplored (19). Our results shed light on the possibility of involvement of PCD in causing DWL. However, the existence of PCD in bacteria seems counterintuitive, the main concern being the benefit of maintaining genes that function to mediate self-destruction of a unicellular organism. PCD may have an altruistic role under conditions of stress. The primary response of a cell, under stressful conditions, is induction of DNA repair mechanisms. However, if the damage is insurmountable, with the cost of repair exceeding the cost of building a new cell, ‘a point of no return’ is reached and PCD is the last resort option adopted by the cell. The demise of some cells could promote the survival of its siblings (37). We hypothesize that upon phage infection majority of the cells undergo lysis, but a sub-population undergoes PCD. The small population of surviving cells, depends on the nutrients released by the dead cells, and may eventually become a nucleus for a renewed population. To the best of our knowledge, this is the first study shedding light on the occurrence of PCD in mycobacteria upon phage infection.

In order to unravel the genetic mechanism underlying phage induced mycobacterial PCD, we investigated the involvement of one of its key regulators, the TA module. We specifically focused on TA loci because of their ubiquitous presence in bacterial genomes and their increasingly observed roles under stressful conditions (38). Several studies conducted on the involvement of the mazEF system in induction of PCD have revealed both pro-survival and pro-death functions (39–41). It seems to play a ‘Janus role’ in determining the fate of a bacterial cell. In our investigation, the TA system genes were found to be universally down regulated upon phage infection.

Collectively our results provide a detailed mechanistic insight into the phage induced mycobacterial cell death. Our study with ‘fluorophages’ is the beginning of a model system for studying the dynamics of phage-host interaction. The appearance of different hallmarks of a PCD pathway in phage infected cells opens an interesting avenue for future research. The potential of this pathway can be tapped by targeting it for future development of a new class of antimicrobials for the treatment of TB.

## Author Contributions

FC was instrumental in carrying out the designed experiments and contributed the text and figures in the manuscript. RS provided assistance in performing immunofluorescence microscopy. MD conducted transmission electron microscopy. SD analyzed the data, conceived the idea and approved the final version of the manuscript.

## Acknowledgement

We thank Prabir Halder for his technical assistance and Ranjan K. Dutta for FACS technical support.

